# Delivery of a chemically modified noncoding RNA domain improves dystrophic myotube function

**DOI:** 10.1101/2025.01.17.633605

**Authors:** Zeinabou Niasse-Sy, Bo Zhao, Ajda Lenardič, Huyen Thuc Tran Luong, Ori Bar-Nur, Johan Auwerx, Martin Wohlwend

## Abstract

Fast twitch, type II muscle fibers are particularly prone to degradation in skeletal muscle pathologies, such as sarcopenia and muscular dystrophies. We previously showed that endogenous activation of the exercise-induced long noncoding RNA CYTOR promotes fast-twitch myogenesis. In the present study, we identify an independent pro-myogenic element within human CYTOR and optimize its RNA delivery. In human primary myoblasts exogenous, vector-based CYTOR^exon^ ^2^ recapitulates the effect of full-length CYTOR by enhancing fast-twitch myogenic differentiation. Furthermore, chemically modified CYTOR^exon^ ^2^ RNA^ΨU^ (N1-me-PseudoU, 7-methyl guanosine 5’Cap, polyA tail) enhanced RNA stability and reduced the immunogenic response to CYTOR^exon^ ^2^ RNA. We demonstrate that viral- or chemically optimized RNA-mediated CYTOR^exon^ ^2^ administration enhances the commitment towards myogenic maturation in Duchenne muscular dystrophy-derived primary myoblasts, induced myogenic progenitor cells and mouse embryonic stem cells. Furthermore, chemically optimized CYTOR^exon^ ^2^ improves key disease characteristics in dystrophic myotubes, including calcium handling and mitochondrial bioenergetics. In summary, our findings identify CYTOR exon 2 as the pro-myogenic domain of CYTOR that can be delivered in a disease context using chemical modifications. This is of particular importance given the susceptibility of type II muscle fibers in different muscle pathologies such as aging and dystrophies, and the reported oncogenic effect of CYTOR exon 1. Our study, therefore, highlights the potential of identifying functional domains in noncoding RNAs. Delivery, or targeting of such RNA domains could constitute next-generation RNA therapeutics.

## INTRODUCTION

Skeletal muscle is an essential organ in humans, playing key roles in locomotion, energy expenditure and metabolic homeostasis. Skeletal muscle is composed of heterogeneous muscle fibers that are broadly classified into two types. Type 1 muscle fibers are slow-twitch, mitochondria-rich, fatigue-resistant and hence, display slow contractile forces. Type 2 fibers are larger, fast-energy producing muscle fibers with rapid force-generating capacity. Notably, type 2 fibers are affected in various disorders such as Duchenne muscular dystrophy (DMD) where type 2 fibers are the first reported muscle fibers that are degenerated(1–3). Type 2 muscle fibers are also affected in other myopathies, including myotonic dystrophy 2(4) and facioscapulohumeral muscular dystrophy(5,6), where they exhibit signs of atrophy. Additionally, sarcopenia and cachexia lead to the deterioration of type 2 skeletal muscle fibers(7–12). Therefore, promotion of type 2 fibers provide an interesting therapeutic avenue for the treatment of these skeletal muscle disorders.

Long noncoding RNAs (lncRNA) have recently emerged as a promising class of biomolecules modulating muscle physiology, amongst many other roles across organs and organisms. Traditionally, they were challenging to target therapeutically but recent advances in RNA technology allow efficient delivery of RNA molecules(13). Recent studies have demonstrated that lncRNAs are involved in the regulation of gene expression in skeletal myogenesis, and that their aberrant expression is linked to multiple muscle disorders(14–16). For example, linc-MYH regulates MuSC proliferation by altering the composition of the INO80 chromatin remodeler complex(17). By stabilizing MYH1B, the lncRNA FKBP1C was shown to promote slow-twitch differentiation(18) whereas SMARCD3-OT1 induced hypertrophy by promoting a fast-twitch muscle fiber phenotype in chicken, which might be relevant to increase meat yields(19). For human therapeutic relevance, however, functional conservation of such lncRNAs in humans remains to be demonstrated. Furthermore, noncoding RNA delivery needs to be optimized in terms of molecule length, stability and immunogenicity.

Our recent findings unveiled the exercise-responsive lncRNA CYTOR in human skeletal muscle, which promotes type 2 muscle fiber formation and maintenance during muscular aging(20). Moreover, employing endogenous overexpression of CYTOR using CRISPRa facilitated differentiation of muscle progenitor cells into mature myotubes, concurrently safeguarding the integrity of fast-twitch, type II muscle fibers in the context of muscle aging. This discovery underscores the significant therapeutic promise of CYTOR to treat muscular disorders characterized by the decline of type 2 fibers. However, despite the promising pro-myogenic effect of CYTOR overexpression, CYTOR was shown to be highly expressed in a variety of tumors including gastric cancer(21,22), colon cancer(23), lung adenocarcinoma(24,25), breast cancer(26,27) and reported to increase tumorigenesis, including invasion, metastasis, malignant proliferation, and glycolysis(28). Overexpression of CYTOR is closely related to clinicopathological characteristics, such as tumor stage, lymph node metastasis/infiltration, and poor prognosis of tumor patients. The dual “moonlighting” effect of CYTOR in myogenesis and oncogenesis suggests that CYTOR has either cell-type specific functions, or contains different functional domains that serve different roles in different cell types. Dissecting these two opposing effects could have important clinical implications to avoid unwanted side effects, such as cancer.

In recent years, the concept of functional domains in lncRNAs has emerged. In mice, a molecule mimicking the functional region of the lncRNA HULC (nt.183-216) was found to be a promising therapy for phenylketonuria(29). Similarly, using the functional domain of Nron lncRNA (nt.3037-3391), a key suppressor of bone resorption, could potentially be a treatment for osteoporosis with reduced side effects compared to the full length Nron lncRNA(30). A recent preprint (available on: https://www.biorxiv.org/content/10.1101/2024.07.23.604857v1) also identifies an ultra-conserved functional domain in the lncRNA CRNDE, which pairs with 32S pre-rRNA to recruit eIF6 to the pre-60S particle. Thus, for lncRNAs with moonlighting roles, it might be beneficial to identify functional domains as different domains might regulate different functions, which might confine side-effects.

Given the dual role of CYTOR in oncogenesis and myogenesis, here we investigated the possibility of functional elements within CYTOR. Gene expression profiling combined with immunocytochemistry suggest that lentiviral overexpression of CYTOR exon 2 but not exon 1 recapitulates full-length CYTOR’s effect on promoting skeletal myogenesis. Optimization of CYTOR^exon^ ^2^ RNA delivery in human cells reveals enhanced RNA kinetics and reduced skeletal muscle inflammatory response using CYTOR^exon^ ^2,m1Ψ^ compared to polyadenylated/5’ capped CYTOR^exon^ ^2^ or naked CYTOR^exon^ ^2^ delivery. Finally, we show that viral- or pulsed RNA delivery of optimized CYTOR^exon^ ^2,^m1Ψ is sufficient to induce fast-twitch maturation in primary muscle cells from Duchenne muscular dystrophy patients and commit induced myogenic progenitors and mouse embryonic stem cells to muscle maturation.

## RESULTS

### CYTOR exon 2 overexpression recapitulates full-length CYTOR overexpression

Among the various isoforms of CYTOR, the transcript ENST00000331944.10 emerges as the most abundantly expressed isoform across multiple human tissues, comprising two exons consisting of 217 nucleotides (exon 1) and 312 nucleotides (exon 2) (**Figure S1A**). We chemically probed the secondary in vitro and in cellulo structure of the main CYTOR RNA isoform using SHAPE-MaP(31), which suggested that the two exons might fold into independent domains, particularly in the cellular context (**Figure S1B-C**). While exon 1 appeared less structured in vitro and in cellulo, exon 2 contains stable structural elements, as shown by their presence in both in vitro and in cellulo predicted structures (**Figure 1A-C and Figure S1D**). Given the structural differences between CYTOR exon 1 and exon 2, the modular nature of RNA(32), and the fact that CYTOR exon 1 has previously been linked to oncogenesis(33), we hypothesized that the two main exons of CYTOR might exert different effects on muscle cell maturation when expressed in isolation.

**Figure 1.**
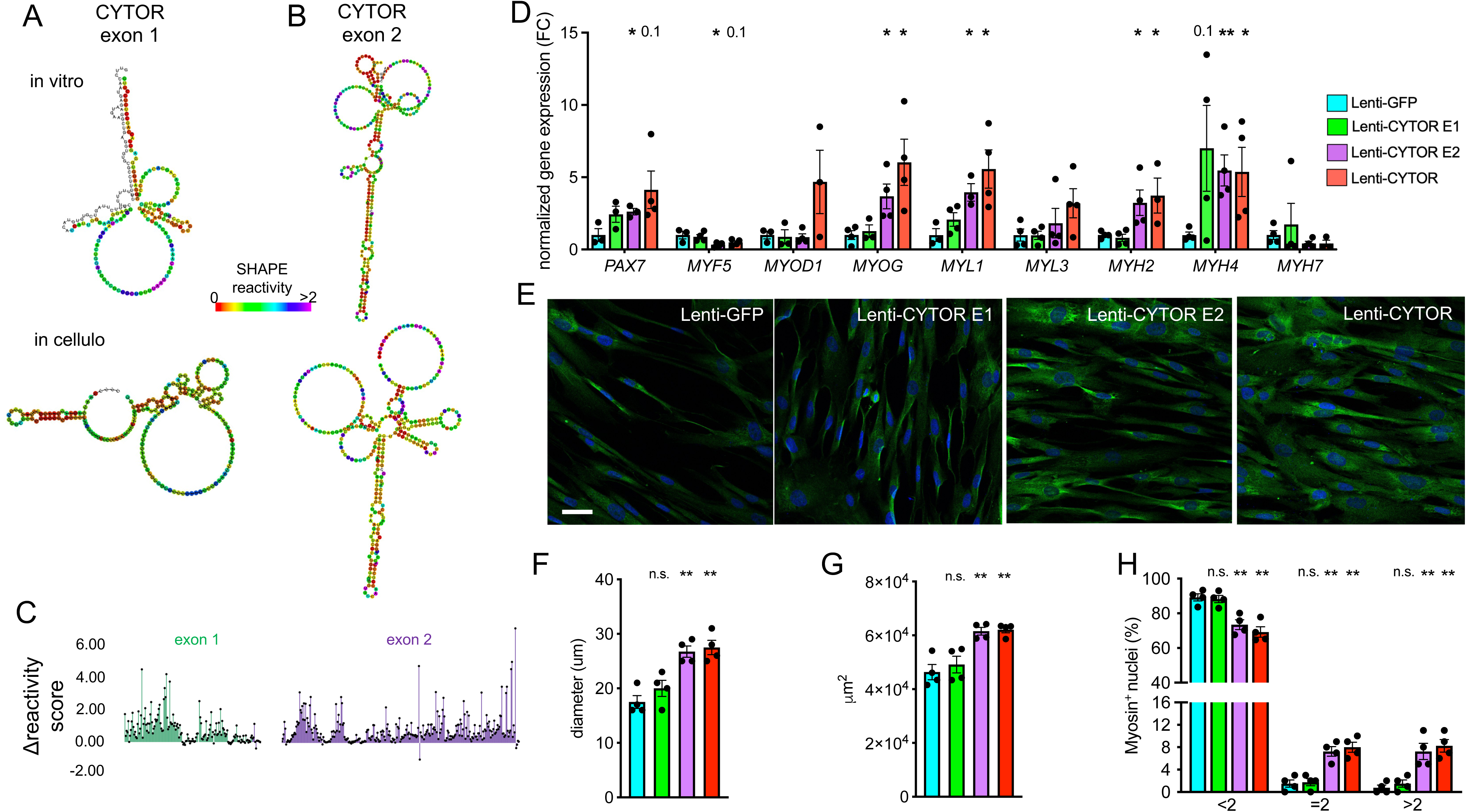
Lentivirus-mediated stable exogenous overexpression of human CYTOR exons in human muscle cells. **(A-B)** SHAPE-directed RNA secondary structure prediction in vitro and in cellulo by selective 2’-hydroxyl acylation analyzed by primer extension and mutational profiling (SHAPE-MaP) of (**A**) human CYTOR exon 1 and (**B**) CYTOR exon 2 in K562 cells. **(C)** Delta SHAPE reactivity scores from (**A-B**) calculated from in vitro and in cellulo SHAPE reactivities. **(D)** Normalized gene expression in differentiating human primary myoblasts transduced with lentivirus capable to express GFP, CYTOR^exon^ ^1^, CYTOR^exon^ ^2^ or full length CYTOR. **(E)** Immunocytochemistry of differentiating human primary myoblasts transduced with lentivirus expressing GFP, CYTOR^exon^ ^1^, CYTOR^exon^ ^2^ or full length CYTOR. **(F)** Quantification of myotube diameter in human primary myoblasts transduced with lentivirus expressing GFP, CYTOR^exon^ ^1^, CYTOR^exon^ ^2^ or full length CYTOR **(G)** Quantification of surface area covered by myotubes in human primary myoblasts transduced with lentivirus expressing GFP, CYTOR^exon^ ^1^, CYTOR^exon^ ^2^ or full length CYTOR. **(H)** Quantification of number of nuclei per myotube in human primary myoblasts transduced with lentivirus expressing GFP, CYTOR^exon^ ^1^, CYTOR^exon^ ^2^ or full length CYTOR. Data: mean ± SEM. *P < 0.05, **P < 0.01.

To assess a potential involvement of CYTOR exon 1 or CYTOR exon 2 in myogenic differentiation, we packaged and transduced lentiviruses containing human CYTOR exon 1, CYTOR exon 2 and full length CYTOR (see **Table S1** for RNA sequences) into primary human skeletal muscle myoblasts obtained from apparent healthy donors. Compared to a GFP control expression construct, exogenous overexpression of full length CYTOR or CYTOR exon 2 for 10 days increased transcripts indicative of myogenesis (*MYOG*; p<0.05) and fast-twitch myotubes (*MYL1*, *MYH2*, *MYH4*; p<0.05) (**Figure 1D**). An increase in MYL1 protein level was confirmed by immunocytochemistry (**Figure 1E**), and quantification of myotube stainings revealed an increase in myotube diameter, overall myotube area and multinucleation of differentiating human muscle cells by overexpressing full length CYTOR or CYTOR exon 2 but not CYTOR exon 1 (**Figure 1F-H**). The finding that exogenous overexpression of full length CYTOR promotes fast-twitch myogenesis is in line with endogenous overexpression of human CYTOR from its genomic locus(20), suggesting that CYTOR might act in trans. In summary, these findings nominate CYTOR exon 2 as the myogenic element of the lncRNA CYTOR, thereby reducing the length to deliver the myogenic element of CYTOR to 312nt.

### Chemical modification of CYTOR exon 2 RNA improves stability and immunogenicity

Virus mediated delivery of transgenes in humans is promising but currently requires high virus titers (>1E14 vg/kg) to achieve clinically relevant expression levels, which can lead to severe immunogenic reactions(34,35). On the other hand, direct RNA delivery is becoming a good alternative to transgene delivery. Reducing the length of an RNA molecule has been shown to increase stability and reduce an immune response(36). Even though we were able to reduce CYTOR length to 312nt, we aimed to further optimize CYTOR delivery by chemically modifying CYTOR RNA nucleotides. To improve the CYTOR exon 2 RNA immunogenic profile and the noncoding RNA stability of exon 2, we considered 5’caps (m7G) and 3’ polyadenylation (polyA) as they have been shown to protect RNAs from exonucleases and help RNA mimic endogenous mRNA, thus reducing the likelihood of triggering unwanted immune responses(37,38). In addition, we took advantage of N1-methylated pseudouridine (m1Ψ), which has been shown to reduce RNA sensing by the cellular innate immune system(39) and increase RNA stability, probably by stabilizing RNA structure(40).

We then synthesized CYTOR exon 2 RNA without any modifications and compared its stability to 5’ capped (m7G) and polyA CYTOR^exon^ ^2^, and 5’ capped (m7G), polyA CYTOR^exon^ ^2,m1Ψ^ in human cells over the time course of 24 hours (**Figure 2A**). As expected, CYTOR exon 2 RNA levels increased robustly up to 10 hours but then normalized again for the remaining 14 hours. While 5’capped, polyA CYTOR^exon^ ^2^ and 5’capped, polyA CYTOR^exon^ ^2,m1Ψ^ showed higher initial levels compared to naked CYTOR exon 2 RNA delivery, there was no apparent difference between the two modified RNA conditions during the first 12h. However, at the 24h timepoint CYTOR^exon^ ^2,m1Ψ^ was more abundant compared to either 5’capped and polyA CYTOR^exon^ ^2^ or naked CYTOR exon 2 RNA delivery, and still 40-fold more abundant than endogenous CYTOR exon 2 RNA levels at the 0h timepoint (**Figure 2F**). To test whether the improved in-cell stability of 5’capped/polyA CYTOR^exon^ ^2,m1Ψ^ RNA observed in cells was due to inherent properties of the RNA modifications, or whether cellular context was needed, we monitored RNA in-solution stability in cell medium without cells for 24h at 37C and compared to plasmid DNA. Plasmid DNA encoding CYTOR exon 2 degraded little (<10%) over 24h. As expected, RNA degraded significantly more compared to DNA (≈ 50%) but there was no difference between different CYTOR^exon^ ^2^ RNA modifications (**Figure S2A**). These results suggest that polyA, m7G and m1Ψ improved CYTOR^exon^ ^2^ RNA stability, particularly when delivered to cells.

**Figure 2.**
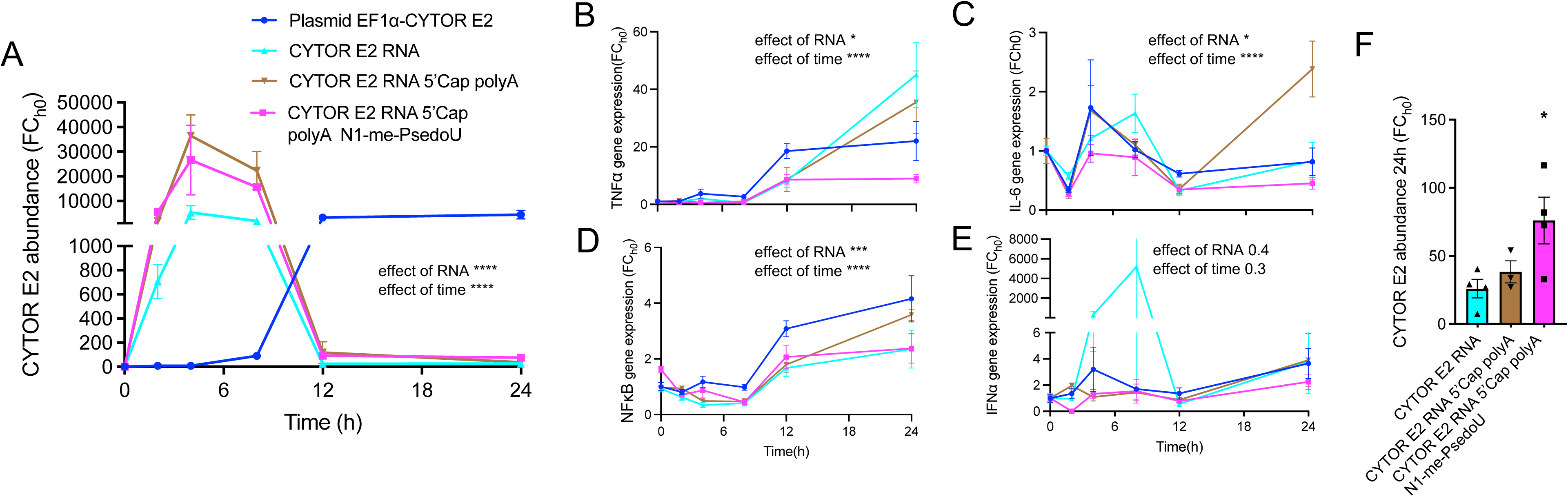
Optimization of RNA-based CYTOR exon 2 delivery. **(A)** Normalized *CYTOR* expression at different time points in HEK293 cells after modified and unmodified CYTOR exon 2 RNA or DNA plasmid delivery. (**B-E)** Time course of normalized gene expression of genes related to the innate immune response (*TNFα*, *IL-6*) and genes induced by pattern recognition receptor activation (*NFκB*, *IFNα*) upon treatment with modified and unmodified CYTOR exon 2 RNA or DNA plasmid delivery. **(F)** *CYTOR* RNA abundance from (**A**) at the 24h time point. Data: mean ± SEM. *P < 0.05, **P < 0.01.

RNA is recognized by pattern recognition receptors (PPRs), hence a commonly reported obstacle to RNA delivery is the strong immune response that is triggered by RNA administration(37). Therefore, we next compared the immunogenic reaction to our different CYTOR exon 2 RNA delivery molecules in human cells. Overall, polyA CYTOR^exon^ ^2,m1Ψ^ with m7G chemical modifications attenuated activation of the innate immune response markers *TNFα* and *IL-6* (**Figure 2B-C**), and showed reduced activation of *NFkB* and *IFNα*, which are typically activated by PPRs upon detection of foreign RNA (**Figure 2D-E**). Altogether, these data suggest that polyadenylated CYTOR^exon^ ^2^ RNA with m7G and m1Ψ chemical modifications displays enhanced RNA stability and an attenuated immunogenic profile in cells.

### CYTOR^exon^ ^2,**Ψ**U^ administration promotes myogenic differentiation in dystrophic myoblasts

Even though chemical modifications of CYTOR^exon^ ^2^ RNA reduced immunogenicity and improved RNA stability, after 12 hours there is still notable less CYTOR^exon^ ^2^ RNA available compared to CYTOR exon 2 RNA derived from virus vectors (**Figure 2A**). Therefore, we next tested whether short exposure with two pulses of our chemically optimized CYTOR^exon^ ^2^ RNA is sufficient to promote myogenic differentiation in human muscle cells. We chose to test CYTOR^exon^ ^2,ΨU^ RNA in myoblasts isolated from Duchenne muscular dystrophy (DMD) patients because these patients have impaired type II muscle fibers(1–3), and we found ≈46% reduced CYTOR levels in myoblasts from DMD patients compared to myoblasts from healthy controls (**Figure 3A**). Two pulses of CYTOR^exon^ ^2,m1Ψ^ RNA were administered in the time course of 4 days and we performed expression and immunohistochemical analyses at day 10 post initiating treatment (**Figure 3B**). Compared to a scramble control RNA sequence, CYTOR^exon^ ^2,m1Ψ^ increased expression of markers indicative of myogenesis and type II muscle cell maturation (**Figure 3C**). Increased MYL1 protein was confirmed by immunocytochemistry (**Figure 3D**), and quantification of emerging myotubes revealed increased myotube diameter, overall myotube area and multinucleation of differentiating human muscle cells treated with chemically modified CYTOR^exon^ ^2,m1Ψ^ RNA compared to a chemically modified scramble control RNA (**Figure 3E-G**). These findings suggest that CYTOR exon 2 can indeed be delivered as RNA and that CYTOR exon 2 RNA is sufficient to promote fast-twitch myogenesis, similar to vector-based CYTOR overexpression.

**Figure 3.**
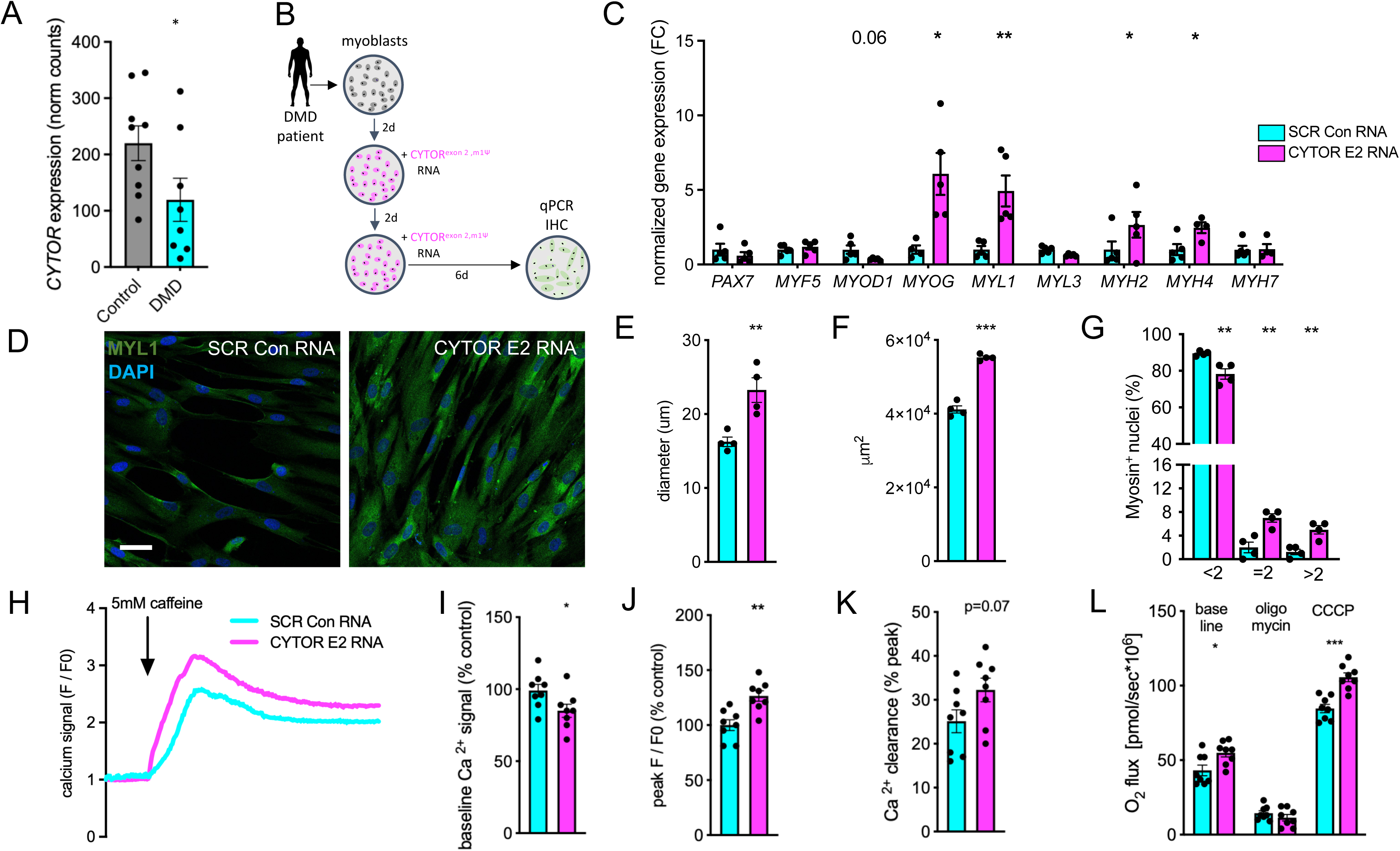
Myogenic effect of optimized RNA gene therapy using the exon 2 myogenic CYTOR element in dystrophic human primary muscle cells. (**A**) Normalized CYTOR RNA levels in primary muscle cells from healthy controls compared to Duchenne muscular dystrophy patients (E-MTAB-8321). (**B**) Schematic showing culturing and treatment regimen of human primary myoblasts from Duchenne muscular dystrophy patients (DMD) with CYTOR exon 2. (**C**) Normalized gene expression after chemically modified CYTOR^exon^ ^2^ RNA administration in differentiating human primary myoblasts isolated from DMD. (**D)** Immunocytochemistry of differentiating dystrophic human primary muscle cells transfected with chemically modified CYTOR^exon^ ^2^ RNA. (**E-G**) Quantification of myotube diameter (**E**), surface area covered by myotubes (**F**), and number of nuclei per myotube in dystrophic human primary muscle cells treated with chemically modified CYTOR^exon^ ^2^ RNA. (**H**) Representative traces of cytosolic, baseline-normalized Ca^2+^ transients in dystrophic myotubes treated with CYTOR exon 2 upon 5mM caffeine stimulation (SR Ca^2+^ store). (**I**) Raw baseline Ca^2+^ signal in dystrophic myotubes treated with CYTOR exon 2 RNA or a scramble control RNA as percentage of control. (**J**) Ca^2+^ amplitude (SR Ca^2+^ store) upon 5mM caffeine stimulation as percentage of scramble control. (**K**) Ca^2+^ clearance as calculated from percentage decrease from the Ca^2+^ peak amplitude. (**L**) Mitochondrial respiration in CYTOR exon 2 and control RNA treated dystrophic muscle cells at baseline and upon addition of oligomycin (1uM) and CCCP (2.5uM). Data: mean ± SEM. *P < 0.05, **P < 0.01, ***P<0.001.

### CYTOR^exon^ ^2,**Ψ**U^ improves disease hallmarks in Duchenne muscular dystrophy derived muscle cells

Duchenne muscular dystrophy (DMD) skeletal muscle is characterized by dysregulated calcium handling(41,42) and impaired mitochondrial bioenergetics(43,44). Therefore, we next tested whether CYTOR exon 2 RNA delivery could modify these key DMD disease characteristics. We first loaded CYTOR^exon^ ^2,m1Ψ^ or scramble control RNA treated myotubes (**Figure 3B**) with the calcium dye Fluo-4 AM and then traced calcium signals from myotubes at baseline, and upon addition of caffeine to release calcium from the sarcoplasmic reticulum stores (**Figure 3H**). We found that CYTOR^exon^ ^2,m1Ψ^ reduced intracellular calcium levels (**Figure 3I**) and increased sarcoplasmic reticulum calcium stores in dystrophic myotubes, as shown by increased peak calcium amplitudes (**Figure 3H and Figure 3J**). After calcium stores were released, there was a trend for improved calcium clearance (**Figure 3K**). Overall, these findings suggest that CYTOR exon 2 enhanced the capacity to store and release calcium in myotubes derived from DMD patients.

We next studied mitochondrial function in dystrophic muscle cells treated with CYTOR exon 2 or a scramble control RNA using high-resolution respirometry in an oxygraph. Results revealed increased baseline oxygen flux in muscle cells treated with CYTOR^exon^ ^2,m1Ψ^ (**Figure 3L**). While there was no difference in mitochondrial respiration driven by oligomycin-induced proton leak, titration with the mitochondrial uncoupler carbonyl cyanide 3-chlorophenylhydrazone (CCCP) revealed improved respiratory capacity in DMD muscle cells treated with CYTOR^exon^ ^2,m1Ψ^ (**Figure 3L**). Elevated levels of pro-inflammatory cytokines such as TNFα and IL-6 have been reported in DMD(45–47) and might therefore, constitute a muscle-cell intrinsic source of tissue inflammation. Notably, measuring expression of these cytokines in dystrophic myotubes from DMD patients showed reduced levels of *TNFα* and *IL-6* upon CYTOR^exon 2,m1Ψ^ treatment (**Figure S2B**). Dystrophic muscle fibers undergo apoptosis as a consequence of dysfunctional dystrophin. Therefore, we next measured muscle cell viability and apoptosis in healthy and dystrophic myotubes treated with CYTOR exon 2. Indeed, CYTOR^exon^ ^2,m1Ψ^ improved dystrophic myotube viability, while simultaneously reducing muscle cell apoptosis as shown by Terminal deoxynucleotidyl transferase dUTP nick end labeling (TUNEL) (**Figure S2C**). In summary, these data show the potential of CYTOR exon 2 RNA to improve key disease hallmarks of DMD, including calcium homeostasis, mitochondrial function, pro-inflammatory cytokine release and muscle cell viability.

### Vector or RNA-based delivery of CYTOR exon 2 enhances early muscle lineage commitment

We next aimed to investigate whether CYTOR exon 2 could promote myogenic differentiation in other muscle progenitor models. Human CYTOR was shown to elicit muscular benefits in *C.elegans*(20), suggesting that the functional (myogenic) effect of human CYTOR exon 2 might be conserved across species. Therefore, we assessed the myogenic potential of human CYTOR exon 2 in two mouse progenitor cell models. Induced myogenic progenitors cells (iMPCs), which represent a fibroblast-derived and expendable mouse muscle progenitor cell model(48). We used plasmid DNA to overexpress CYTOR exon 2, or transfected CYTOR^exon^ ^2,m1Ψ^ RNA twice into differentiating iMPCs and harvested them 10 days later to measure markers of muscle cell maturation. Results showed an overall increase in transcripts indicative of muscle cell differentiation, particularly fast-twitch myosins with no apparent difference between vector-based or RNA-based CYTOR exon 2 delivery (**Figure 4A-B**). We then went one step further back in cell fate commitment and delivered CYTOR exon 2 using a vector, or CYTOR^exon^ ^2,m1Ψ^ RNA to differentiating mouse embryonic stem cells (mESCs)(49). Similar to our findings in iMPCs, CYTOR exon 2 delivery into mESCs increased transcripts associated with myogenic differentiation (**Figure 4C-D**). These results suggest that exon 2 of human CYTOR shows cross-species commitment towards a muscular fate by promoting fast-twitch myogenesis. In summary, the present study shows that exon 2 of the long noncoding RNA CYTOR is sufficient to promote myogenesis (**Figure 5**). This finding is particularly important considering the oncogenic effect of CYTOR exon 1.

**Figure 4.**
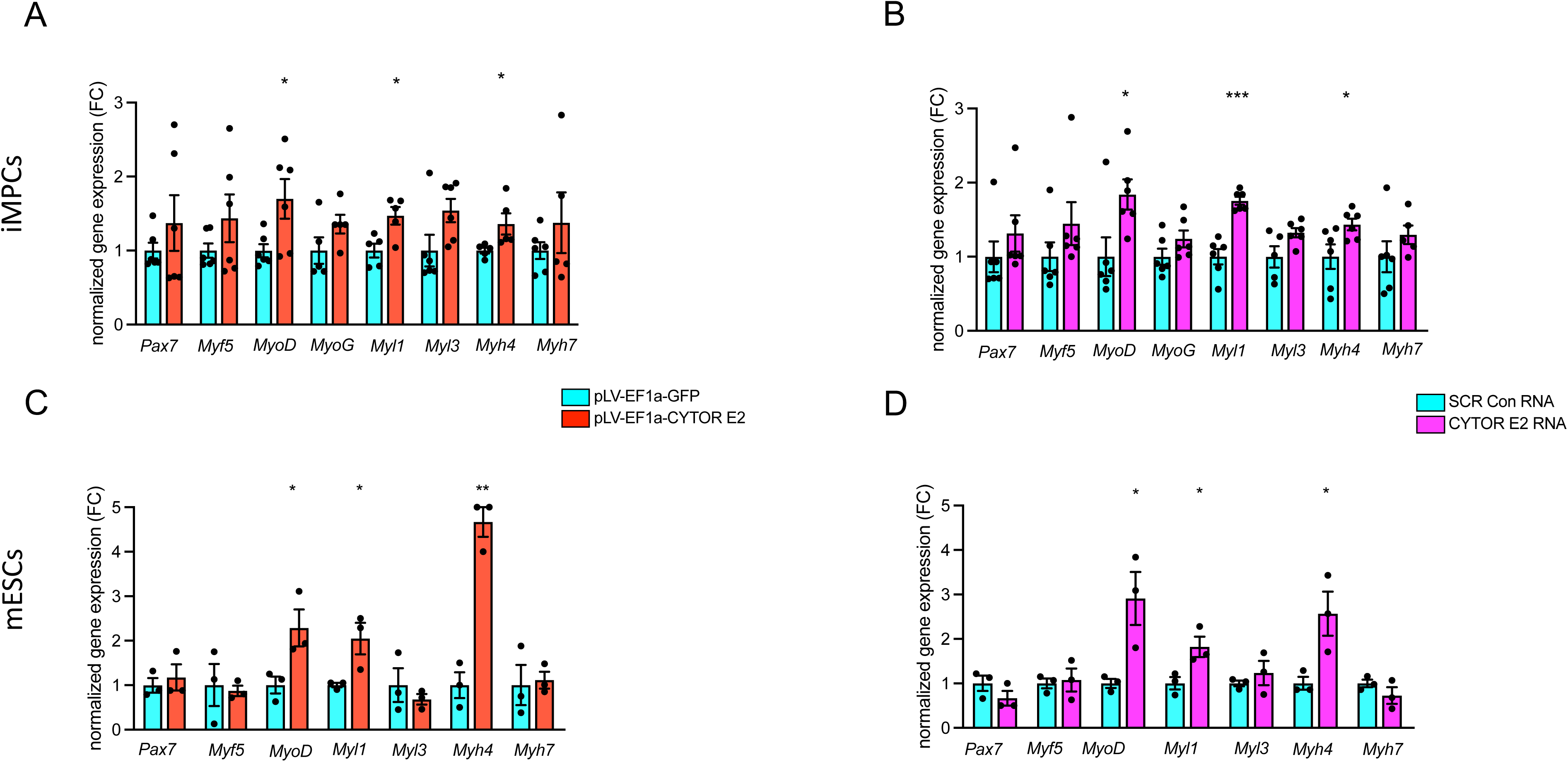
Forced expression of human CYTOR exon 2 in mouse induced myogenic progenitor and embryonic stem cells. (**A**) Normalized gene expression in mouse embryonic stem cells transduced with lentivirus expressing a GFP control construct or CYTOR^exon^ ^2^. (**B)** Normalized gene expression in mouse embryonic stem cells after transfection of chemically modified CYTOR^exon^ ^2,ΨU^ RNA. **(C)** Normalized gene expression in induced myogenic progenitors transduced with lentivirus expressing a GFP control construct or CYTOR^exon^ ^2^. (**D**) Normalized gene expression in induced myogenic progenitors after transfection of CYTOR^exon^ ^2,ΨU^ RNA. Data: mean ± SEM. *P < 0.05, **P < 0.01.

**Figure 5.**
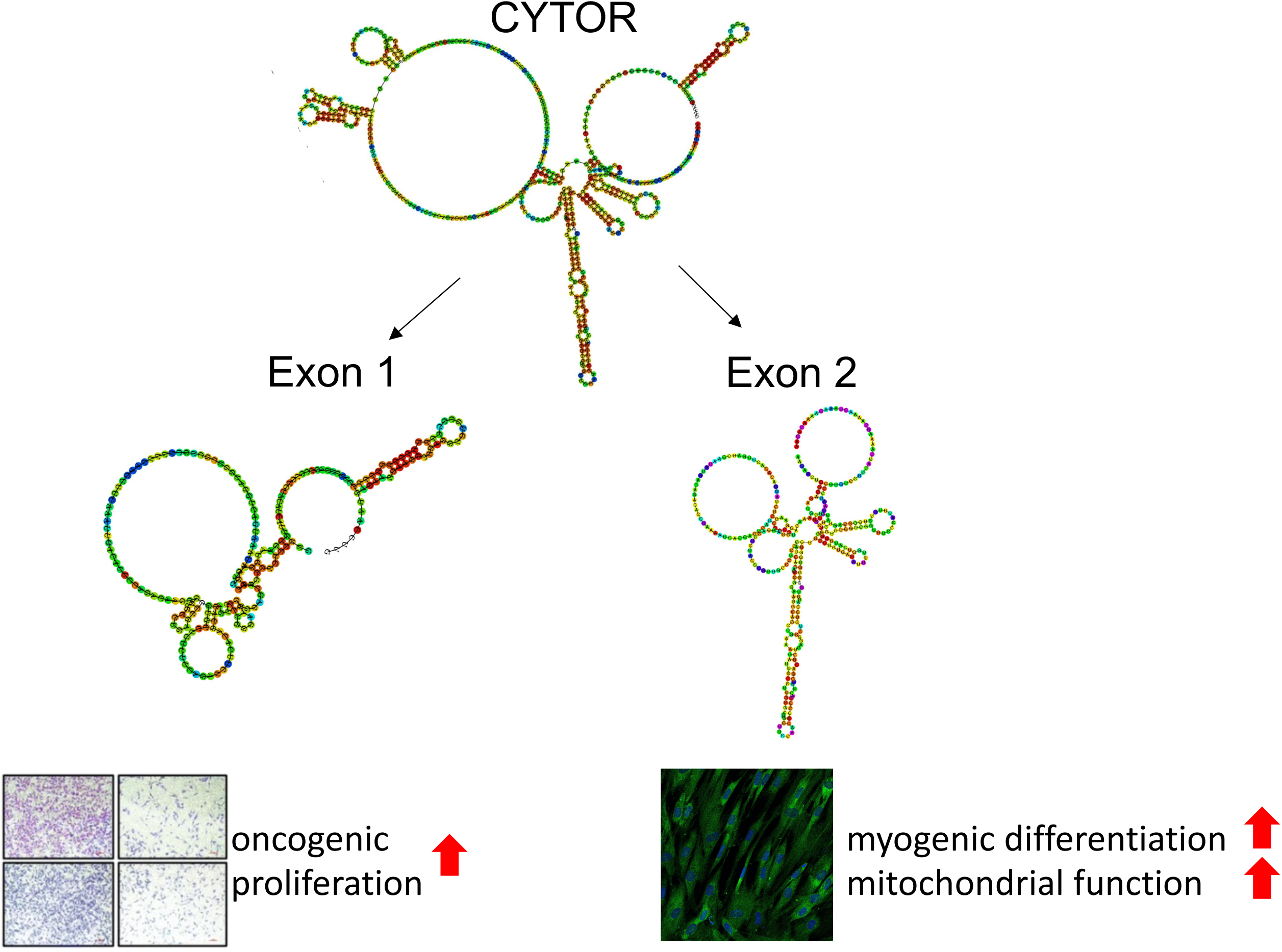
Summary of functional elements of CYTOR driving myogenesis and oncogenesis. Schematic overview over the different effects exon 1 and exon 2 exert. CYTOR exon 1 has been shown to drive cancer cell proliferation (33). In this study we identify CYTOR exon 2 to promote muscle cell differentiation.

## DISCUSSION

The major findings of the present study show that **I)** CYTOR exon 2 recapitulates the pro-myogenic effect of full-length CYTOR, **II**) N1-me-PseudoU (m1Ψ), m7G 5’cap and poly-A modifications render CYTOR exon 2 RNA more resilient to degradation and reduce its immunogenic response, **III**) CYTOR exon 2 RNA delivery enhances fast-twitch myogenesis in pluripotent stem cells, muscle progenitors and dystrophic primary myoblasts in mouse and human. Therefore, reducing the length of CYTOR RNA coupled with chemical modification of the functional CYTOR exon 2 RNA element might constitute an attractive way of delivering the functional element of CYTOR to promote myogenesis for therapeutic applications.

Noncoding RNA (ncRNA) therapeutics are promising due to the cell-type specific expression profiles of noncoding RNAs. However, ncRNAs that regulate key cellular pathways, such as CYTOR, might be hijacked by cancers. Therefore, systemic delivery of such ncRNAs might no longer be a safe option unless subdomains in these noncoding RNAs can be identified that exert different effects in cancer vs normal tissues. Multiple studies recently reported pro-oncogenic effects driven by CYTOR in breast, stomach, colon and hepatocellular cancers. Importantly, this oncogenic activity was demonstrated to be specifically exerted by exon 1 of CYTOR(33), suggesting the existence of functional elements within CYTOR RNA. The concept of functional domains in lncRNAs is supported by the recent discovery of a 20nt stretch within the lncRNA HULC which recapitulated the full-length effect of HULC on phenylalanine tolerance(29,50). In addition, delivery of the full-length lncRNA Nron not just increased bone resorption but also caused splenomegaly, likely due to a strong immune response, whereas delivery of just the conserved functional motif of Nron effectively reversed bone loss without side effects(30). Moreover, bulk deletions and CRISPR-mediated targeting of NEAT1 revealed domains that affect paraspeckle formation and cancer growth(51). Therefore, previous studies suggest the existence of functional domains in lncRNAs. In the present study we identify exon 2 in CYTOR driving myogenic differentiation which could explain the moonlighting role of CYTOR in both oncogenesis and myogenesis. Given the oncogenic effect of CYTOR exon 1, our findings suggest that identification of a functional domain in a noncoding RNA might further increase spatial specificity depending on where functional ncRNA domains are active. Previous studies employed organ-targeting ligands for tissue-specific delivery of lncRNA mimetics(29,52). Combining domain-specific targeting with tissue-specific ligands, or decorated lipid nanoparticles, might therefore, further enhance cell type-specific action of ncRNAs in the future.

Considering ncRNA functional domains as potential therapeutic targets has several benefits. First, reducing target length could facilitate delivery of long noncoding genes that exceed packaging limitations of current gene therapy technologies (≈ 4.8kb). Second, in the case of RNA delivery, shorter cargoes reduce the immunogenetic response to the RNA. By identifying exon 2 as the myogenic driver of CYTOR, were able to reduce the length of CYTOR needed to promote myogenesis to 312nt. From a translational perspective, reducing CYTOR RNA length to be delivered by ≈45% could not only reduce RNA manufacturing costs (synthesis and purification) and thus, increase scalability, but also enhance CYTOR RNA stability and facilitate delivery when packaged in lipid nanoparticles, for example. Even though we did not test the immune response to full length CYTOR, longer RNA constructs have been shown to elicit larger immune responses(36). Therefore, reducing length of CYTOR coupled with chemical modifications might act in synergy to reduce adverse side effects (immune response) and improve ncRNA bioavailability. We improved the immunogenic profile of CYTOR exon 2 RNA delivery using m1Ψ (renowned by its successful use on COVID-19 mRNA vaccines) and m7G modifications. Our study underlines the effectiveness of m1Ψ chemical modifications on reducing noncoding RNA immunogenicity, particularly when delivering a functional element. Besides improved immune profiles with these chemical modifications m1Ψ CYTOR exon 2 delivery additionally showed a 40 fold increased abundance 24h after delivery compared to unmodified CYTOR exon 2, thereby increasing the biological availability of the functional CYTOR element.

Intriguingly, only two pulsed doses of CYTOR exon 2 RNA were sufficient to drive muscle progenitor differentiation. This finding suggests that the improved muscle maturation we observed might be the results of a self-reinforcing cycle, once myoblasts are pushed over the "activation energy" threshold needed for differentiation. In general, short exposure to ncRNAs could therefore, be harnessed for epigenetic/cellular priming and reprogramming. Direct RNA delivery has thus-far mostly been used for vaccines and to delivery Cas9/gRNAs for gene editing. Here we show that short exposure to a functional ncRNA element was a sufficient epigenetic cue to prime cells for differentiation.

Despite poor conservation of lncRNAs, we previously showed that CYTOR, when expressed in *C. elegans* (no CYTOR orthologue) triggered similar muscular benefits as in mouse and human. Here, we expressed human CYTOR exon 2 in mouse muscle cell models and find that human CYTOR also exerts comparable effects as was observed with endogenous overexpression of the mouse Cytor orthologue(20). Cross-species effects of CYTOR exon 2 suggests that the effect of the pro-myogenic exon 2 element is also conserved. CYTOR’s effect across different species could be explained by CYTOR acting in trans. This is possible as we observed comparable myogenic effects of human CYTOR whether it was overexpressed locally(20) or exogenously (this paper).

Our work has some limitations. We showed that exon 2 partially recapitulated the effect of full-length CYTOR but we cannot completely rule out an early effect on muscle differentiation by exon 1. Moreover, even though we observed overall similar phenotypes between human and murine models upon treatment with CYTOR exon 2, there were small differences in gene expression profiles between human and murine models. These differences might be related to species-specific myogenic gene programs or the stemness of our progenitor cell models. While our work suggests putative benefits of CYTOR exon 2 treatment in patient-derived samples, experiments were performed in in vitro models. Even though cell autonomous effects are well captured in human cell systems, these models lack the complexity of in vivo animal models. More preclinical animal work is needed to establish type 2 fiber promotion as a viable strategy for DMD, and other dystrophies.

In summary, we find human CYTOR exon 2 to exert pro-myogenic effects in mouse and human cell cultures, mimicking full length CYTOR. Biologically, these findings intriguingly suggest the existence of functional elements in the lncRNA CYTOR. From a therapeutic perspective, targeting exon 2 using vector strategies or optimized RNA delivery avoids oncogenicity as was shown with exon 1 of CYTOR. Moreover, chemical modifications to improve noncoding RNA stability and reduce immunogenicity, together with shortening the length of CYTOR might improve its translational potential in a clinical setting. Overall, our study highlights the potential of identifying functional elements in ncRNAs as delivery or targeting of such RNA domains could constitute next-generation RNA therapeutics.

## METHODS

### *In vitro* mouse and human studies

#### Primary muscle cell culture and cell transfection

Primary human skeletal muscle cells were obtained from Lonza (SkMC, #CC-2561) and primary human myoblasts from Duchenne muscular dystrophy patients were provided by Hospices Civils de Lyon and cultured in growth medium consisting of DMEM/F12 (Gibco, 10565018), 10 % Fetal Bovine Serum (Gibco, 10270-106) and 100 U/mL penicillin and 100 mg/mL streptomycin (Gibco, 15140-122). Cell culture medium was changed to 2% Fetal Bovin Serum to support myoblast differentiation. To control for experimental variation, the same batch of FBS was used throughout the study. Time of experiments varied for the different assays performed in this study. Cell transfections were done using TransIT-X2 (Mirus) according to the manufacturer’s protocol with a 3:1 ratio of transfection agent to DNA. All cells were maintained at 37 °C with 5% CO2. Experiments were performed in the period 2019-2025.

#### RNA isolation and real-time qPCR

Cells were homogenized, and RNA isolated using the RNeasy Mini kit (Qiagen, 74106). Reverse transcription was performed with the High-Capacity RNA-to-cDNA Kit (4387406 Thermofisher scientific) as shown previously(53). Gene expression was measured by quantitative reverse transcription PCR (qPCR) using LightCycler 480 SYBR Green I Master (50-720-3180 Roche). Quantitative polymerase chain reaction (PCR) results were calculated relative to the mean of the housekeeping gene *GAPDH*. The average of two technical replicates was used for each biological data point. Primer sets for qPCR analyses are shown in the **Table S2**. To measure CYTOR expression in samples from Duchenne muscular dystrophy patients we analyzed normalized counts from a publicly available RNA sequencing dataset as previously shown(54). In brief, we accessed E-MTAB-8321 and extracted the day 0 myoblast RNA sequencing counts of CYTOR in healthy controls and Duchenne muscular dystrophy patients. We next performed the Grubbs’ test, also called the ESD method (extreme studentized deviate), to determine whether there is a significant outlier from the rest. Replicate 3 of sample M197 was excluded from the analysis as it was a significant outlier (p<0.05).

#### Secondary RNA structure probing

Selective 2′-Hydroxyl Acylation analyzed by Primer Extension and Mutational Profiling (SHAPE-MAP) was used as previously described (31) to analyze RNA structure at nucleotide resolution. In brief, RNA in vitro or in cellulo were treated with 2-methylnicotinic acid imidazolide (NAI) that reacts with flexible nucleotides at the 2’-hydroxyl group, marking them without disrupting base pairing (Eclipsebio, California, United States). After RNA isolation (in cellulo), RNA was then reverse transcribed into cDNA, during which modified nucleotides introduce mutations or stop reverse transcription. The cDNA libraries were then sequenced, and we performed mutational calling to calculate the mutational rates of NAI- and DMSO control treated samples. We then computed an overall reactivity profile by subtracting the two mutational rates. SHAPE reactivities were then normalized. The difference between in vitro and in cellulo SHAPE reactivities was calculated as delta reactivity and used to analyze differences in vitro and in cellulo structures. RNA secondary structures were predicted using a previously published folding algorithm and visualized using the Vienna RNA folding tool (55,56).

#### Induced myogenic progenitor cell culture

Mouse embryonic fibroblasts (MEFs) Induced myogenic progenitor cells (iMPCs) were cultured as previously described (48). Reprogramming of low-passage MEFs into iMPCs was done with medium containing KnockOut-DMEM (ThermoFisher Scientific, 10829-018) supplemented with 10% FBS (HyClone SH30396.03), 10% KnockOut Serum Replacement (ThermoFisher Scientific, 10828028), 1% GlutaMAX (35050061), 1% non-essential amino acids (ThermoFisher Scientific, 11140050), 1% penicillin-streptomycin (ThermoFisher, 15140122), 0.1% β-mercaptoethanol (ThermoFisher Scientific, 21985-023) and 10ng/ml basic FGF (R&D 233-FB). Forskolin (Sigma Aldrich F6886) and RepSox (Sigma-Aldrich, R0158) were added at 5μM each. 3μM of the GSK3β inhibitor CHIR99021 (Tocris) was used. Doxycycline (Sigma-Aldrich, D9891) was added at a final concentration of 2μg/ml to induce expression of Myod that was previously transduced as a dox-inducible expression system into MEFs using lentivirus. Expanded bulk cultures as well as iMPC clones were cultured in iMPC medium without dox and supplemented with Forskolin (5μM), RepSox (5μM), and GSK3β inhibitor (3μM).

#### Mouse embryonic stem cell culture and muscle differentiation

Mouse embryonic stem cells were cultured in KO-DMEM (Thermofisher 10829018) with 1% GlutaMAX-I (Thermo Fisher 35050061), 1% non-essential amino acids (Thermo Fisher 11140050), 1% Pen-Strep (Thermo Fisher 15140122), 0.1% 2-Mecaptoethanol (Thermo Fisher 21985023), 15% FBS (Thermo Fisher 10270-106) and 1000U/mL mouse LIF (PolyGene PG-A1140-0010). Over the time course of 2 weeks mESC were differentiated to muscle cells using a previously published protocol(49). In brief, 4d prior to differentiation, 3uM CHIR99021 and 1uM PD0325901 were added to mESCs in gelatin-coated dishes (0.1-0.2% gelatin). Cells were then resuspended in NK1 medium at 12-17k per cm2 and switched to DF15 CDL medium that contains 0.1uM LDN193189 but no PD0325901 on day 2. On day 4, cells were switched to DK14F1 CDL medium that contains 14% KSR. On day 6, 0.5% DMSO and PD173074 was added at 250nM whereas CHIR99021 was removed. 2 days later 2% horse medium was added and changed every 2 days until day 14 when multinucleated fast MyHC+ myotubes can be observed. Human CYTOR plasmid or RNA was administered at day 2 and day 6 of this differentiation protocol.

#### Calcium transient measurements

Calcium tracing was performed as previously described (42,57). In brief, cultured myotubes were loaded with the Ca^2+^ indicator Fluo-4 AM (5μM; Invitrogen, Basel, Switzerland) diluted in a Krebs Ca^2+^ solution [135.5mM NaCl, 1.2mM MgCl2, 5.9mM KCl, 11.5mM glucose, 11.5mM Hepes, and 1.8mM CaCl2 (pH 7.3)] for 20 min in the incubator and rinsed twice with Krebs solution. Immediately prior to imaging, cells were washed twice with Ca^2+^-free Krebs solution and kept in that solution. Fluo-4 fluorescence traces were monitored using confocal microscopy (Zeiss LSM 5 Live, 40× oil-immersion lens). Excitation wavelength was 488 nm, and the emitted fluorescence was recorded between 495 and 525nm using time-lapse acquisition. After recording basal fluorescence, muscle cells were stimulated with 5mM caffeine (O1728-500, Thermo Fisher Scientific). The Zen software (products/microscopy-software/zenlite/zen-2-lite) was used for the imaging, and analyzed in excel. The raw calcium signal prior to caffeine addition was used to compare baseline calcium levels, whereas calcium peak and clearance were calculated from baseline-normalized calcium levels. The amplitude of Ca^2+^ transients indicative of SR Ca^2+^ stores was calculated by subtracting peak fluorescence from baseline. Ca^2+^ clearance after SR Ca^2+^ release was calculated as the percentage decrease from the peak to the stabilized Ca^2+^ signal.

#### Lentivirus production and transduction

Lentiviruses were produced by cotransfecting HEK293T cells with lenti plasmids expressing GFP or CYTOR EXON 1, EXON 2 or full length (see plasmid list in **Table S3**), the packaging plasmid psPAX2 (addgene #12260) and the envelope plasmid pMD2G (addgene #12259), in a ratio of 4:3:1, respectively. Plasmids were obtained from Vectorbuilder. Virus containing medium was collected at 48h and concentrated using Lenti-X^TM^ Concentrator (Takara). 8μg/mL polybrene (Millipore) for 30min was used to facilitate virus transduction.

#### Mitochondrial respiration

Treated muscle cells were detached from the plates and washed in medium containing 10mM Ca-EGTA buffer, 0.1uM free calcium, 20mM imidazole, 20mM taurine, 50mM 2-(N-morpholino) ethane-sulfonic acid hydrate, 0.5mM dithiothreitol, 6.56mM MgCl_2_, 5.77mM ATP, 15mM phosphocreatine (pH 7.1). To perform high-resolution respirometry, cells were then counted to ensure equal loading across groups, and added to chambers of an O2k oxygraph (Oxygraph-2k; Oroboros Instruments, Innsbruck, Austria) pre-warmed at 37C with MiR05 medium (58,59) containing 110mM sucrose, 60mM K^+^-lactobionate, 0.5mM EGTA, 3mM MgCL2, 20mM taurine, 10mM KH_2_PO_4_, 20mM HEPES and 1g/l bovine serum albumin (pH 7.1) for high-resolution respirometry. Each chamber was used as a biological replicate and chambers were switched during every run. Once the oxygen signal stabilized, baseline was measured for 3 minutes. Then, oligomycin was added at 1uM to inhibit the ATP synthase and thus, measure mitochondrial respiration driven by proton leak. Next, titration (0.5uM steps) of carbonyl cyanide 3-chlorophenylhydrazone (CCCP) to a 2.5uM final concentration uncoupled mitochondria from ATP generation and allowed measurement of uncoupled respiratory capacity. Finally, rotenone (1uM), and antimycin A (1uM) were injected into the chambers to terminate respiration by inhibiting complex I and complex III, respectively and to subtract all measurements from residual, non-mitochondrial, oxygen consumption.

#### RNA production and chemical modification

CYTOR exon 2 RNA was transcribed by co-transcription using NEB’s HiScribe T7 ARCA mRNA kit (with or without tailing) reagent (Vectorbuilder). In short, Anti-Reverse Cap Analog (ARCA, NEB #S1411) using T7 RNA Polymerase was used for the capped CYTOR exon 2 RNA by combining the ARCA/NTP mix, T7 RNA Polymerase Mix and a suitable DNA template (NEB, E2060S). After a brief DNase I treatment to remove the template DNA, capped mRNA is poly(A) tailed using Poly(A) polymerase prior to RNA purification. Chemically modified CYTOR exon 2 mRNA (5-methylcytidine and pseudouridine modified) was synthesized and purified using Silica membranes with two RNeasy steps (TriLink BioTechnologies, San Diego, CA, USA). The same batch of synthesized RNA was used throughout the experiments performed in this study.

#### Cell viability and apoptosis assays

Healthy and dystrophic myotubes were treated with scramble control RNA or chemically optimized CYTOR exon 2 in 96 well plates. Cell viability was measured following the instructions from the Promega CellTiter-Glo luminescence kit (G7571, Promega, United States) using a plate reader (Spark, Tecan, Switzerland). Adding the mixed buffer and substrate to the cells allowed luminescence based quantification of ATP in the cells. Cell apoptosis was measured using a terminal deoxynucleotidyl transferase dUTP Nick End Labeling (TUNEL) assay (4822-96-K, Biotechne). In brief, cells were fixed with 3.7% formaldehyde, permeabilized with Cytonin^TM^ and labeled with Terminal deoxynucleotidyl Transferase (TdT) and a nucleotide mix prior to treating with Strep-HRP to bind labeled DNA. Adding TACS-Sapphire™, a colorimetric substrate, which reacts with Strep-HRP produces a measurable color change, which was then measured and quantified at 450nm.

#### Immunocytochemistry

Lonza muscle cells were cultured on a sterilized coverslip in 6-well plates (Greiner bio-one, CELLSTAR, 657160) were fixed in Fixx solution (Thermo Scientific, 9990244) for 15 min and permeabilized in 0.1% Triton X-100 (Amresco, 0694) solution for 15min at 20°C. Cells were blocked in 3% BSA for 1h at 20°C to avoid unspecific antibody binding and then incubated with primary antibody over night at 4°C with gentle shaking. MyHC was stained using the MYL1 antibody (1:140, Thermofisher, PA5-29635). Antibodies used in this study are shown in Table S4. The next day cells were incubated with secondary antibody (#A-21206 for MYL1) for 1h at 20°C and nuclei were labeled with DAPI. The immunofluorescence images were acquired using either fluorescence or confocal microscopy. Myofusion index was calculated as the ratio of nuclei within myotubes to total nuclei. Myotube diameter was measured for 8 myotubes per image using ImageJ. Myotube area was calculated as the total area covered by myotubes.

#### Statistical analyses

Required sample size was estimated from previous performed experiments with similar readouts and variation in the assays(53,58,42,20). Muscle differentiation and calcium handling experiments were performed blinded. Data analyses for all experiments was performed unblinded. Unpaired student t-tests were applied for comparisons between 2 groups. For experimental conditions in which there were multiple comparisons, a factorial ANOVA with subsequent post hoc analysis was performed. Where appropriate, one-way ANOVA with post hoc analysis or t-tests was performed. All statistical analyses were performed using GraphPad Prism (version 10, San Diego, CA, USA). Results are reported as means ± standard error of the mean. Statistical significance was set to P < 0.05.

## Ethical statement

All patient-derived materials utilized in this study were anonymized to ensure the protection of individual identities and confidentiality. Prior to sample collection, informed consent was obtained from all participants, after comprehensive explanation of the study’s purpose, methods, and potential implications. The use of patient samples was strictly confined to applications in biological research and was conducted in full compliance with applicable ethical guidelines and institutional review board approval 2665.2 (01/03/2019).

## Acknowledgments

We wish to thank the staff of EPFL histology and bioimaging and optics (BIOP) for technical assistance. The work in the JA laboratory was supported by grants from the European Research Council (ERC-AdG-787702), the Swiss National Science Foundation (SNSF 31003A_179435), the Fondation Suisse de Recherche sur les Maladies Musculaires (FSRMM) and the Fondation Marcel Levaillant (190917). MW’s position was supported by Central Norway Regional Health Authority.

## Author contributions

The study was conceived and designed by ZN and MW. Human muscle cell experiments including gene expression profiling and immunocytochemistry were conducted by ZN. *In vitro* work associated with ESCs and iMPCs were performed by AL and supervised by OBN. MW and ZN wrote the manuscript, and all authors gave critical comments on it. MW supervised the work.

## Supplementary Figures

**Figure S1.**
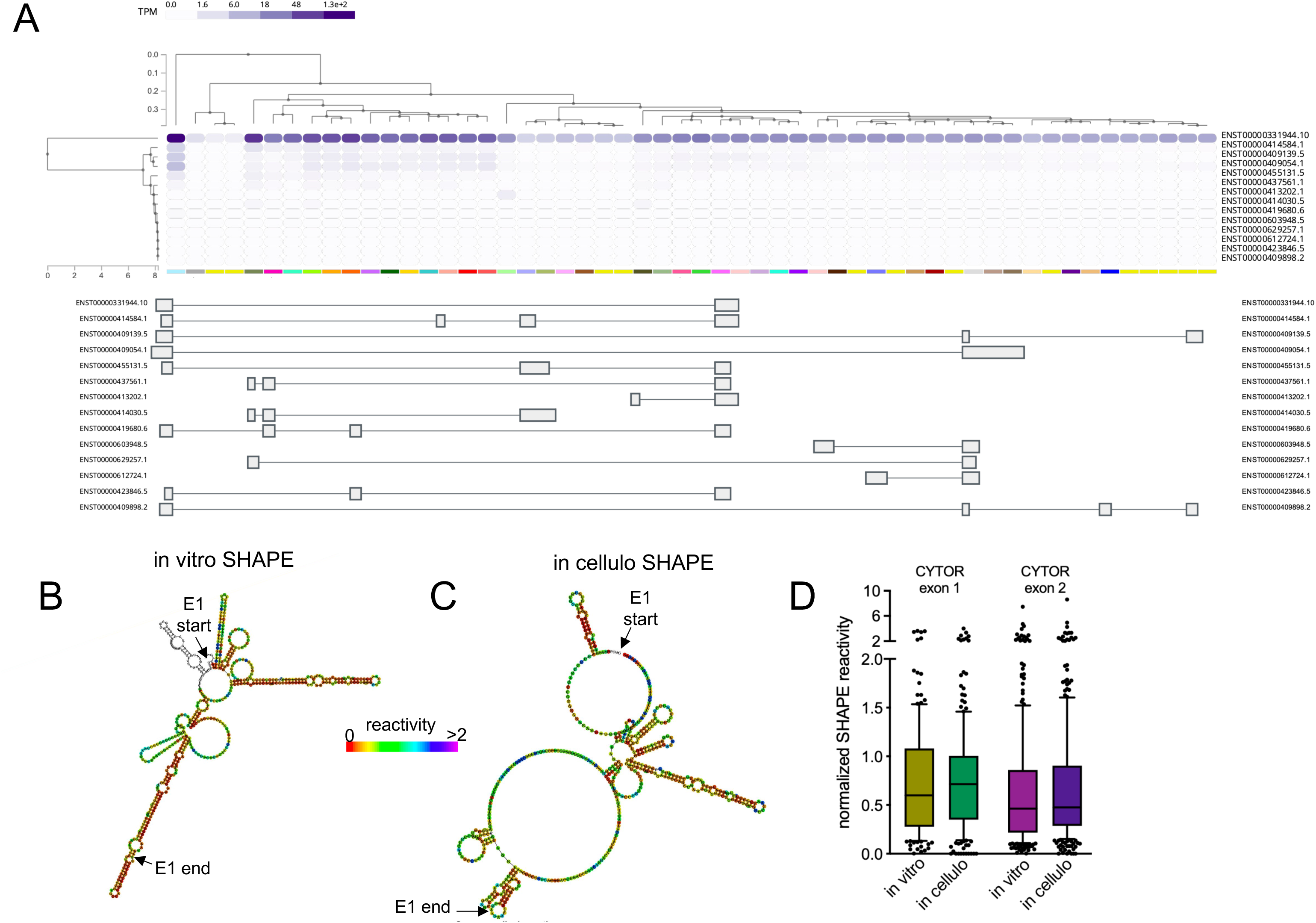
Human CYTOR exon 1 and exon 2. (**A**) CYTOR RNA isoform expressed across 54 human tissues in the Genotype-Tissue Expression (GTEx) portal (accessible on: https://gtexportal.org/home). (**B-C**) SHAPE directed RNA secondary structure prediction of full length CYTOR (**B**) in vitro and (**C**) in cellulo from K562 cells. (**D**) Box plots showing SHAPE reactivity distributions for CYTOR exon 1 and exon 2.

**Figure S2.**
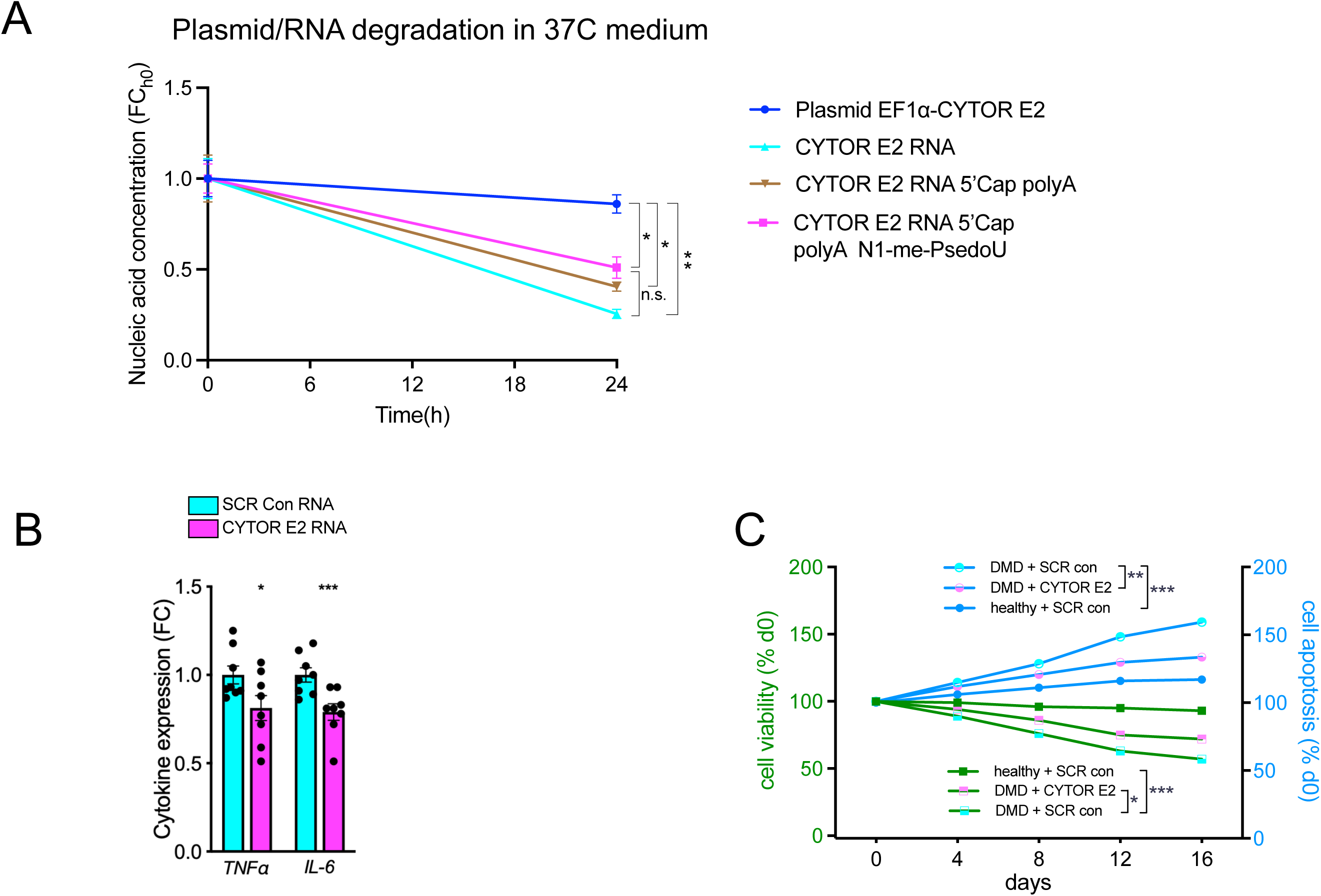
(**A**) Nucleic acid concentration over 24h of CYTOR plasmid or CYTOR exon 2 RNA in standard cell culture medium at 37C. Concentrations were measured with nanodrop. (**B**) Normalized abundance of *TNFα* and *IL-6* in skeletal muscle myotubes from Duchenne muscular dystrophy patients treated with chemically modified CYTOR exon 2 RNA or scramble control RNA (n=8 per group). (**C**) Cell viability and apoptosis in dystrophic myotubes and healthy myotubes treated with scramble control RNA, or chemically optimized CYTOR exon 2 RNA (n=8 per group). Data: mean ± SEM. *P < 0.05, **P < 0.01, ***P<0.001.

**Table S1.**
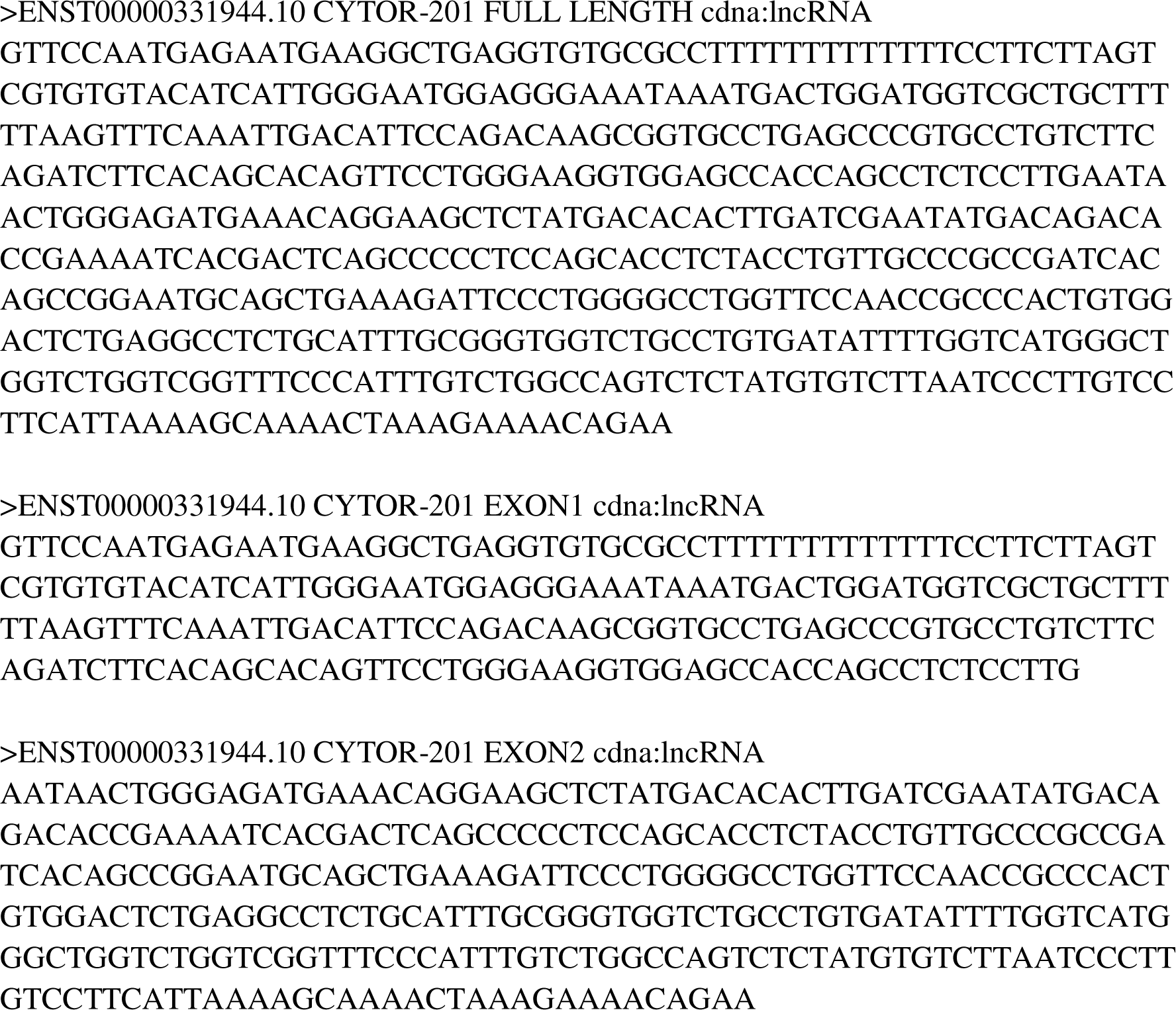
Human CYTOR exon sequences.

**Table S2.**
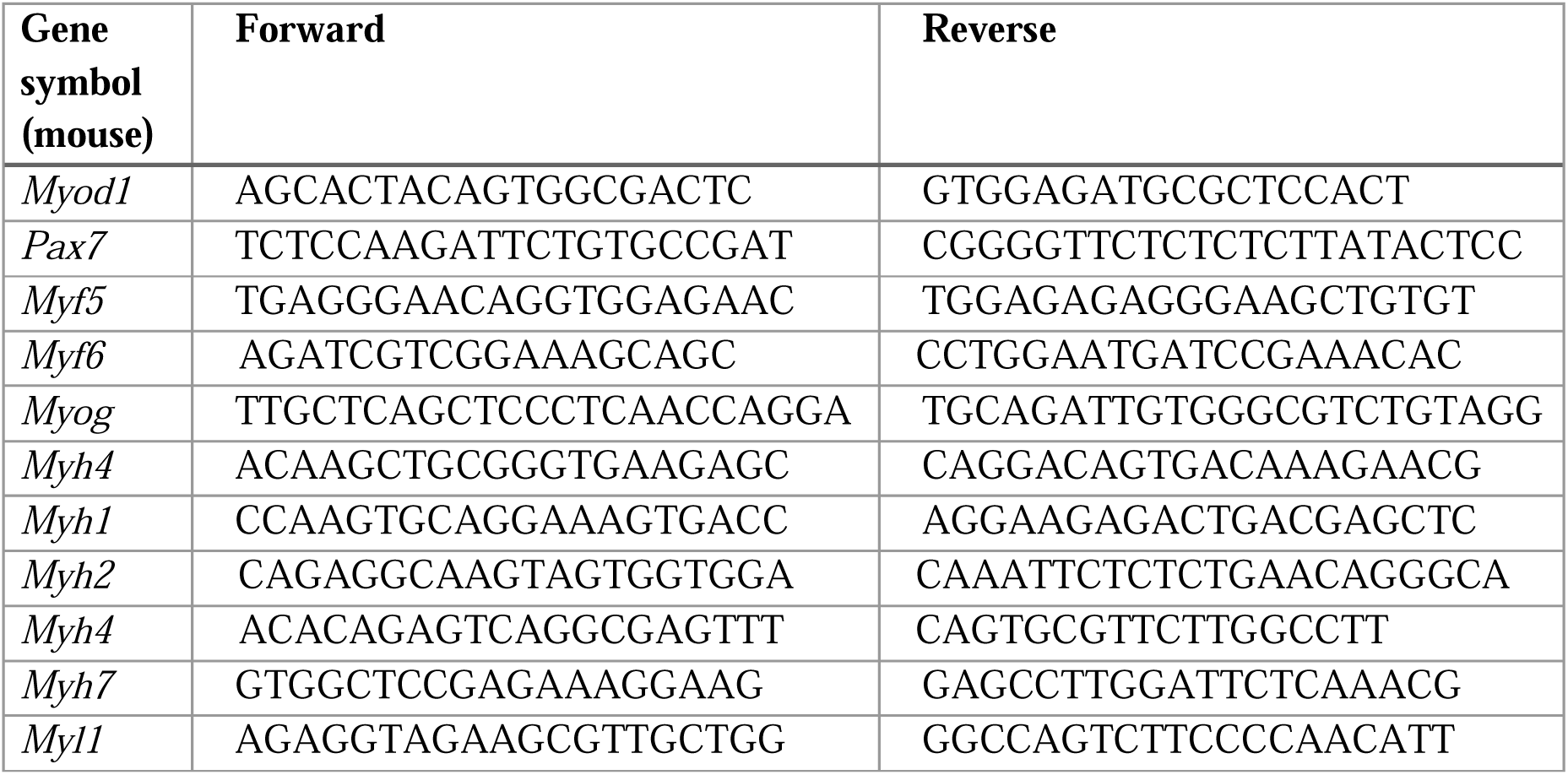
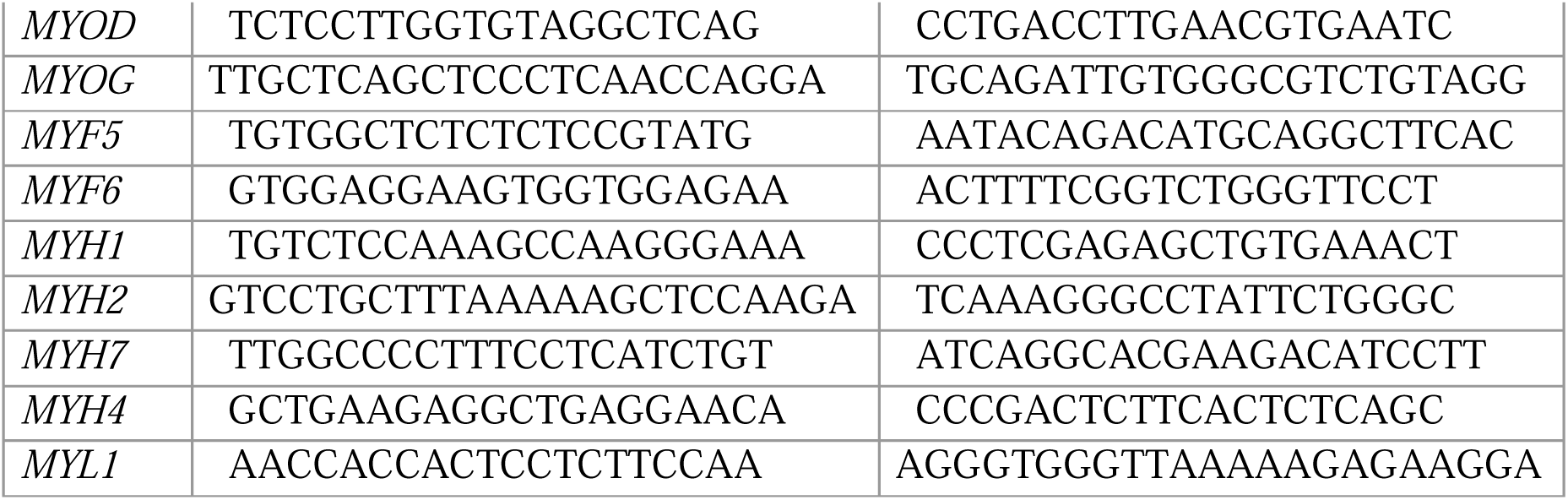
List of mouse and human qPCR primers.

**Table S3.** List of plasmids.

| Plasmid | Reference |
| --- | --- |
| pLV-EF1a-GFP | this paper |
| pLV-EF1a-CYTOR | this paper |
| pLV-EF1a-CYTOR-E1 | this paper |
| pLV-EF1a-CYTOR-E2 | this paper |
| psPAX2 | Addgene # 12260 |
| pMD2G | Addgene # 12259 |

**Table S4.** List of antibodies.

| Antibody | Supplier | Reference # |
| --- | --- | --- |
| Anti-Myosin light chain 1 | Thermofisher | # PA5-29635 |
| Donkey anti-Rabbit IgG secondary antibody | ThermoFisher Scientific | # A-21206 |

